# A novel NGS-compatible Enzymatic Strategy Enables Carryover Contamination Removal and Enhances Sequencing Performance

**DOI:** 10.1101/2025.08.01.668201

**Authors:** Amanda DeLiberto, Laurence Ettwiller

## Abstract

Carryover contamination during DNA amplification can often lead to false positives. Traditional mitigation strategies often include physical separation and/or enzymatic decontamination. However, these methods have limitations, including logistical constraints and polymerase compatibility issues especially for Next Generation Sequencing (NGS). Here, we introduce a novel approach using 7-deaza-dGTP and Fpg for carryover amplicon degradation. When incorporated into library preparation, Fpg degrades 7-deaza-deoxyguanosine amplicons providing carryover protection comparable to the established dUTP/UDG strategy with over 95% carryover contamination removal. Unlike currently available carryover enzyme/dNTP solutions, 7-deaza-deoxyguanosine is compatible with many polymerases and does not impact substitution frequencies during sequencing. Additionally, due to its chemical properties, incorporating 7-deaza-dGTP during amplification improves GC bias during sequencing. In turn, this method is compatible with NGS, supports broader polymerase compatibility, and improves sequencing performance, particularly in AT- and GC-rich regions.

## 1. Introduction

Accurate and contamination-free DNA and RNA amplification is essential for molecular biology applications, including clinical diagnostics, forensic analysis, and next-generation sequencing (NGS). However, one of the major challenges in amplification-based methods is carryover contamination, which can lead to false-positive results and compromised data integrity. This is particularly problematic in low-input or precious samples, where the amount of amplified DNA can greatly exceed the amount of DNA from the starting material. Contaminant DNA from previous amplification reactions can be inadvertently introduced into new reactions, leading to unintended amplification and misinterpretation of results^1^.

Two primary strategies are commonly employed to mitigate carryover contamination in DNA and RNA amplification. The first approach involves implementing stringent physical separation between pre- and post-amplification^2^. However, due to logistical constraints, strict spatial segregation is not always feasible. Another strategy involves enzymatic decontamination of amplicons containing deoxyuridine (dU) in place of deoxythymidine (dT). In this approach, dUTP is substituted for dTTP in the nucleotide mix and incorporated into the amplified DNA during PCR. In the event of cross-contamination in subsequent reactions, the dU-containing amplicons can be selectively degraded by uracil-DNA glycosylase (UDG), which recognizes and excises uracil residues, resulting in the formation of abasic sites^3–6^. During subsequent denaturation at 95°C, these abasic sites lead to strand cleavage, effectively preventing the reamplification of contaminant DNA. Alternatively, strand cleavage can be achieved directly using a mixture of UDG and DNA glycosylase-lyase Endonuclease VIII also known as USER™. Recently, a carryover approach with deoxyinosine (dI)/Endonuclease V was also introduced with compatibility with bisulfite converted amplicons^3^. However, these enzymatic approaches require the use of specific polymerases compatible with dITP/dUTP incorporation and may pose challenges for downstream applications, as dU and dI are recognized as a damaged base by many enzymes. Due to these limitations, enzymatic strategies have rarely been extended to NGS applications.

Here, we introduce a novel strategy using 7-deaza-2’-deoxyguanosine-5’ triphosphate (7-deaza-dGTP) as a substitute for canonical dGTP. We demonstrate that the incorporation of 7-deaza-dGTP into amplicons allows for the selective degradation by formamidopyrimidine-DNA glycosylase (Fpg). We further demonstrate that Fpg selectively and efficiently eliminates 7-deaza-deoxyguanosine–containing amplicons in simulated carryover experiments, offering a robust alternative to the conventional dUTP/UDG system carryover. Importantly and contrasting with existing enzymatic strategies for carryover prevention, this strategy is compatible with standard Illumina sequencing workflows. Furthermore, the use of 7-deaza-dGTP has been shown to improve amplification especially in regions with high GC content^4–11^, and as a result, subsequent DNA sequencing of the 7-deaza-deoxyguanosine-containing fragments provides more even coverage. Overall, this 7-deaza-dGTP/Fpg approach provides an effective and flexible solution for eliminating carryover contamination in nucleic acid amplification and NGS workflows.

## 2. Results

### 2.1 Fpg cleaves 7-deaza-deoxyguanosine in DNA

An earlier study demonstrated that a 7-deaza-deoxyguanosine located in position 15 of a 25 nucleotide long oligonucleotide is recognized by *E. coli* Fpg and cleaved^12^. Other studies have independently demonstrated that DNA polymerases can both incorporate 7-deaza-dGTP and extend from templates containing 7-deaza-deoxyguanosine^4,6,7,9,11^. We therefore hypothesize that the 7-deaza-dGTP/Fpg pair can be used as an alternative to the conventional dUTP/UDG strategy for mitigating carryover contamination that is compatible with NGS sequencing (principle in **Figure 1A**).

To experimentally demonstrate this, we first tested whether the Fpg activity on 7-deaza-deoxyguanosine is retained in double stranded DNA amplified with 7-deaza-dGTP fully replacing dGTP. We observe complete digestion of DNA containing 7-deaza-deoxyguanosine when digested with Fpg, while canonical DNA amplicons remain intact (**Figure 1B**). This result indicates that Fpg recognizes 7-deaza-deoxyguanosine in double stranded DNA independent of context.

**Figure 1:**
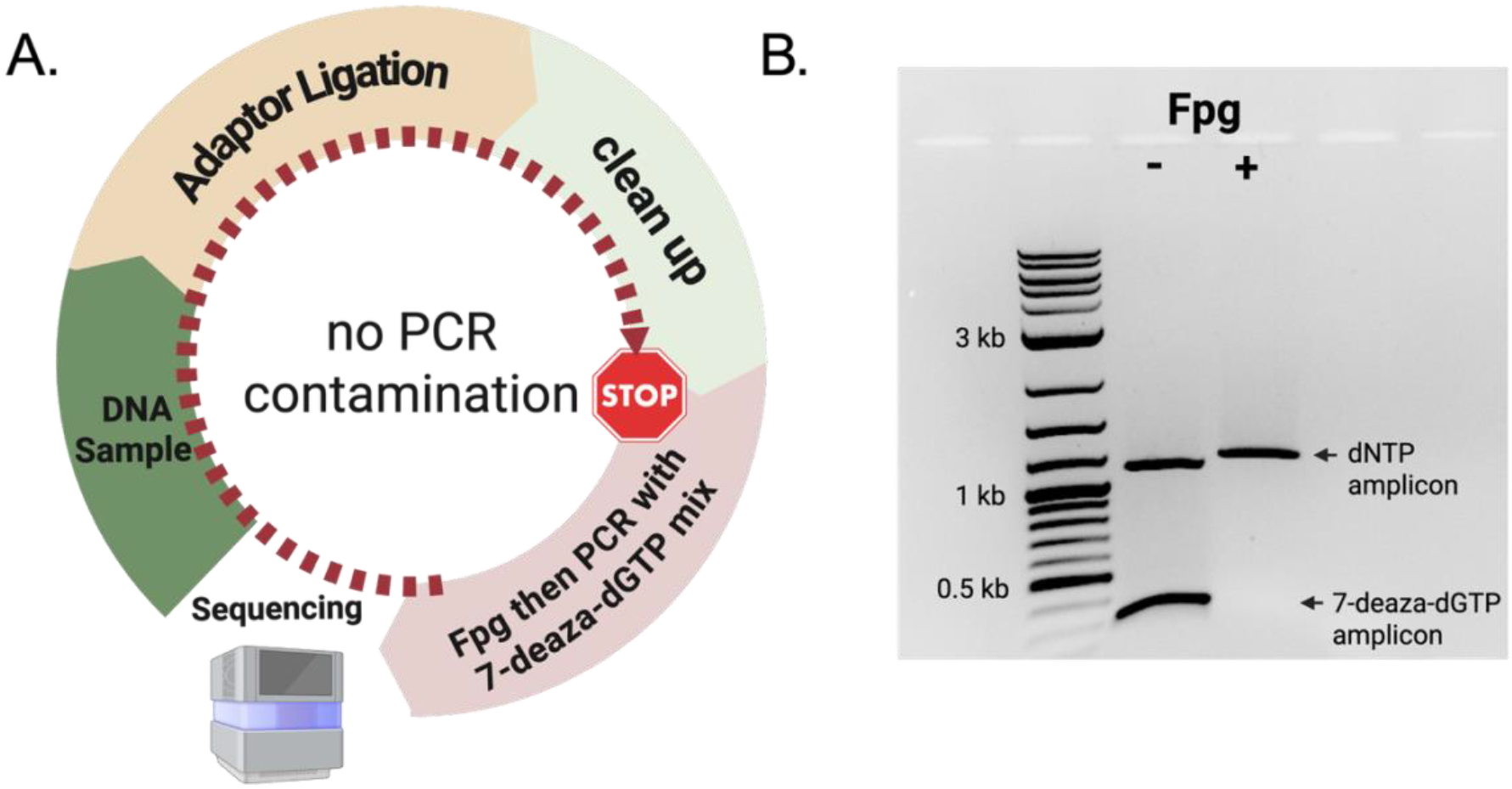
Fpg digests amplicons containing 7-deaza-deoxyguanosine and can be utilized for NGS carryover prevention. **A)** Proposed carryover contamination prevention strategy using Fpg and 7-deaza-dGTP. PCR amplification is performed using a modified nucleotide mix composed of dATP, dCTP, dTTP, and 7-deaza-dGTP. In the event of carryover contamination from previously amplified products, these contaminant amplicons can be selectively cleaved and eliminated through treatment with formamidopyrimidine DNA glycosylase (Fpg) followed by amplification. **B)** A mixture of two Lambda DNA amplicons of distinct sizes—amplified with 7-deaza-dGTP (300 bp) or standard dNTPs (1200 bp), was subjected to enzymatic treatment with Fpg or no enzyme and subsequently analyzed by 1.2% agarose gel.

### 2.2 Use of 7-deaza-deoxyguanosine in NGS

Contrary to the dUTP/UDG strategy, the 7-deaza-dGTP/Fpg strategy does not require specialized Du tolerant polymerases for amplification^4,7,9,11^. Furthermore, incorporation of 7-deaza-dGTP in amplicons has been shown to decrease duplex stability and hence, reduce the stability differences between AT and GC rich regions.^5–7,9^. We therefore hypothesized that the 7-deaza-dGTP/Fpg strategy could be effectively integrated into next-generation sequencing (NGS) workflows without compromising data quality. To evaluate its performance relative to standard NGS workflows and the conventional dUTP/UDG-based carryover prevention method, genomic libraries were prepared from *Escherichia coli* K-12 (50.5% GC), as well as from a mixed genomic sample containing *Staphylococcus epidermidis* (AT-rich, 32% GC) and *Streptomyces coelicolor* (GC-rich, 72% GC). These bacterial species were selected to assess the impact of 7-deaza-deoxyguanosine incorporation on sequencing coverage across a wide range of genomic GC content.

Libraries were amplified using three nucleotide mixes: (1) a modified dNTP mix d[ATC]TP in which the dGTP was fully substituted with 7-deaza-dGTP (7-deaza-dGTP mix), (2) a standard dNTP mix or (3) a modified mix d[AGC]TP in which the dTTP was fully substituted with dUTP (dUTP mix). To accommodate dU containing templates, the dUTP mix libraries were amplified with Q5U, a modified version of the DNA polymerase for which the dU pocket has been modified to avoid stalling at dU^13–15^. Resulting libraries were paired-end sequenced using Illumina and the resulting reads were mapped to a composite reference genome.

#### 2.2.1 7-deaza-deoxyguanosine improves GC bias in NGS

We first evaluated the impact of fully substituting dGTP with 7-deaza-dGTP on insert sizes and sequencing coverage relative to GC content. Libraries amplified with the 7-deaza-dGTP mix have similar size distribution than the other libraries with the notable exception for fragments from *S. epidermidis* where libraries prepared with the standard dNTP mix showed overall shorter inserts (**Supplementary Figure 1A**). 7-deaza-dGTP mix libraries displayed more uniform sequencing coverage across the full GC spectrum compared to those amplified with either the dUTP mix or the standard dNTP mix (**Figure 2, Supplementary Figure 1B**). Notably, genomic regions with extreme GC biases, such as within *S. coelicolor*, or *S. epidermidis*, were often underrepresented or entirely absent in libraries amplified with the standard or dUTP mixes. Conversely, those same regions show improved coverage with libraries amplified with 7-deaza-dGTP mix (**Supplementary Figure 1C**). This observation is consistent with the established use of 7-deaza-dGTP to alleviate secondary structure formation and polymerase stalling during PCR amplification of GC-rich sequences.

**Figure 2:**
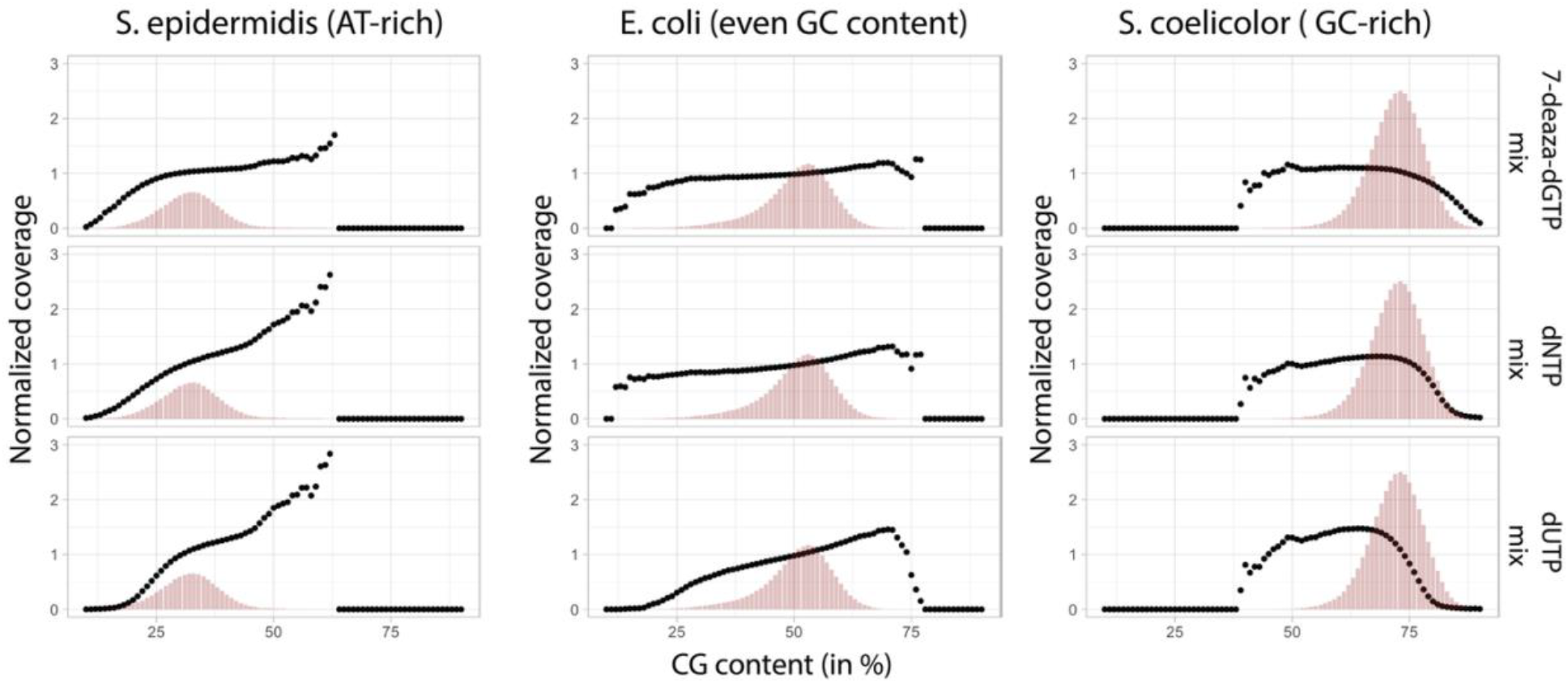
Genome wide GC bias and coverage. Coverage bias plot of the theoretical GC distribution (red bars) and normalized read coverage within 100 bp windows (black) for S. epidermidis (left panels), E. coli (middle panels) and S. coelicolor (right panels) genomes when libraries were either amplified with Q5 (NEB #M0493) using the 7-deaza-dGTP mix (top panels), dNTP mix (center panels) or Q5U with dUTP mix (bottom panels).

**Supplementary Figure 1:**
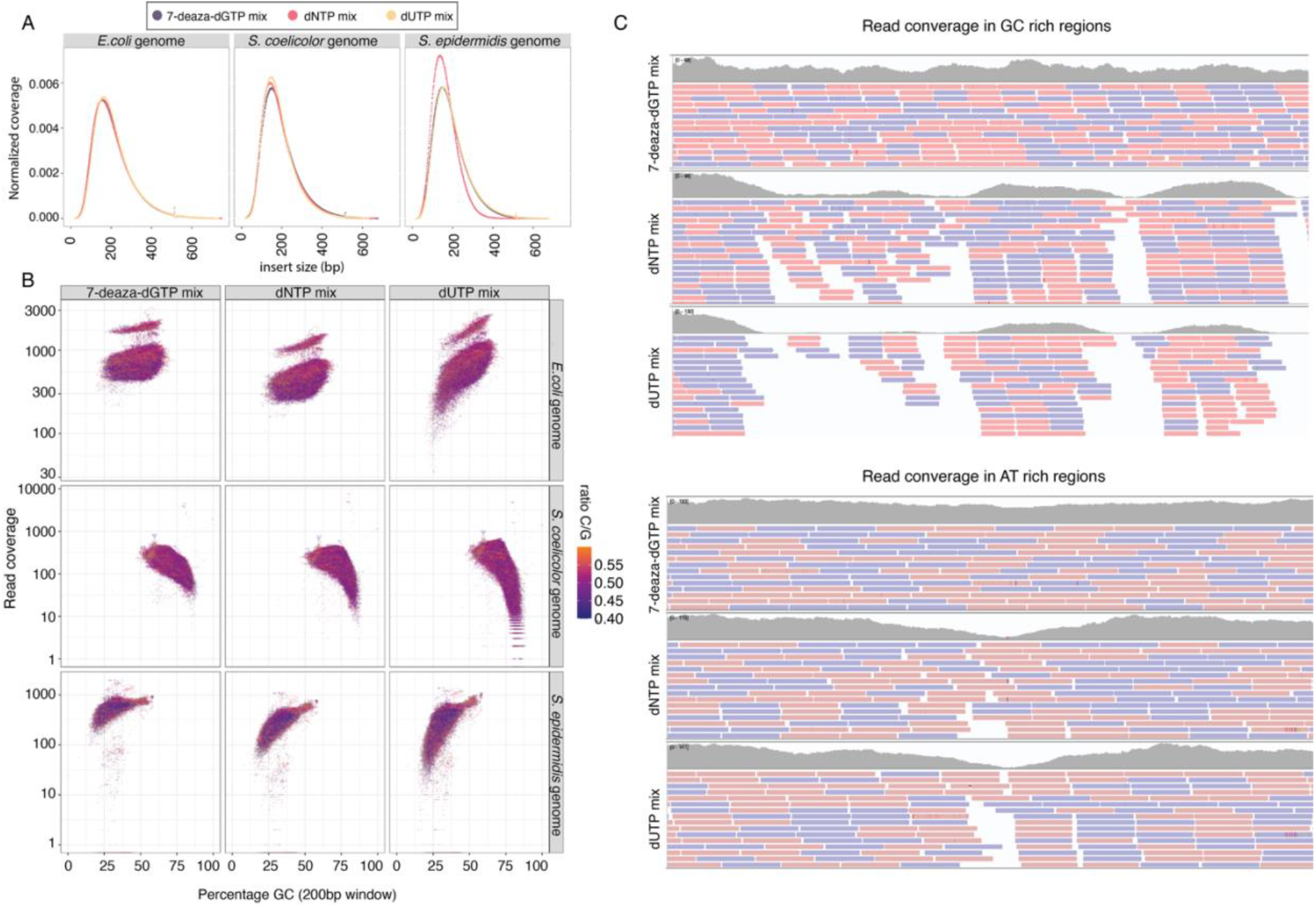
Library size, GC bias and coverage. **A)** Distribution of Insert sizes from paired-end mapping to E. coli genome (left panel), S. coelicolor genome (center panel) and S. epidermidis genome (Right panel). **B)** Read coverage function of the percentage of GC in individual 200 bp sliding windows across S. coelicolor (GC rich), S. epidermidis (AT rich) and E. coli (even GC) with 7-deaza-dGTP mix, dUTP mix, or standard dNTPs. **C)** Integrative Genomics Viewer (IGV) view of read coverage in GC (top) and AT (bottom) rich genomic regions for each dNTP mix.

#### 2.2.2 7-deaza-deoxyguanosine has little or no impact on Sequencing Accuracy

Next, we assessed whether sequencing accuracy metrics in libraries amplified using the 7-deaza-dGTP mix were comparable to those generated with the standard dNTP and dUTP mixes. Overall, substitution frequencies in libraries prepared with the 7-deaza-dGTP mix closely matched those obtained with standard dNTPs (**Figure 3**) with a modest increase in G-to-A substitution frequencies (p =0.013, after Bonferroni correction) using the 7-deaza-dGTP mix compared to the dNTP mixes. Notably, the most significant differences between the 7-deaza-dGTP and standard dNTP mixes were detected for G-to-T (p = 1.96 × 10^−5^, Bonferroni-adjusted) and C-to-A (p = 7.35 × 10^−7^, Bonferroni-adjusted) substitutions, with 7-deaza-dGTP libraries showing significantly lower substitution frequencies relative to the dNTP controls or dUTP libraries.

**Figure 3:**
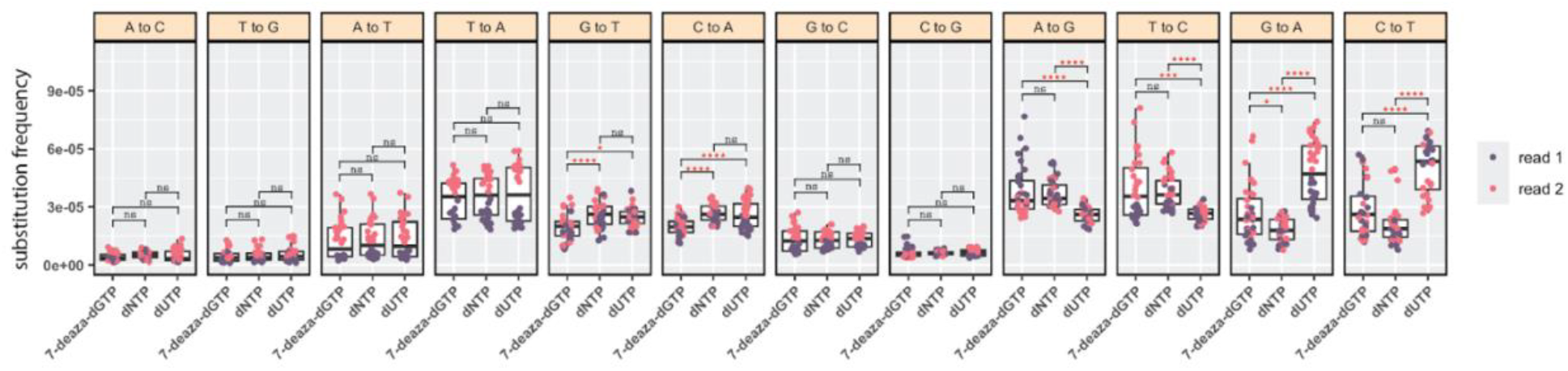
Substitution frequency using different amplification dNTP mixes. For each substitution type, frequencies were calculated separately for paired-end Read 1 (purple) and Read 2 (pink) across all 5′NXN3′ sequence contexts (where N = A, T, C, or G, and X denotes the substitution type). Data are presented for libraries amplified with three different nucleotide mixes: 7-deaza-dGTP mix (left), standard dNTP mix (center), and dUTP mix (right). Statistical comparisons were performed using pairwise t-tests, and p-values were adjusted for multiple testing using the Bonferroni correction method. Significance is denoted (ns: not significant, ^*^< 0.05, ^**^<0.01, ^***^<0.001, ^****^<1e-04)

### 2.3 Fpg effectively eliminates DNA templates containing 7-deaza-deoxyguanosine

To compare the efficiency of carryover contamination removal using Fpg and USER in an NGS context, a pre-constructed library of *S. coelicolor and S. epidermidis* was amplified using standard dNTP mixes. Separately, an *E. coli* library was amplified using either dUTP or 7-deaza-dGTP mixes and subsequently spiked into the amplified *S. coelicolor/S. epidermidis* library at approximately 1:1 ratio. The reciprocal experiment, in which the *S. coelicolor/S. epidermidis* library was spiked into the *E. coli* library to serve as the contaminant, was also performed to control for potential biases related to GC content. The resulting mixes were treated either with Fpg or USER as appropriate. After enzyme treatment, libraries were reamplified using either dUTP or 7-deaza-dGTP mixes as appropriate and sequenced to determine the percent reads mapping to the contaminating library. Upon Fpg treatment we observe almost complete removal of the contaminating 7-deaza-dGTP library **(Figure 4A**,**B)**. This was consistent among replicates and comparable to dUTP/USER protocol (**Figure 4A**,**B**). Overall, we observe over 99% removal of carryover contamination with Fpg/7-deaza-dGTP (**Figure 4C)**.

**Figure 4:**
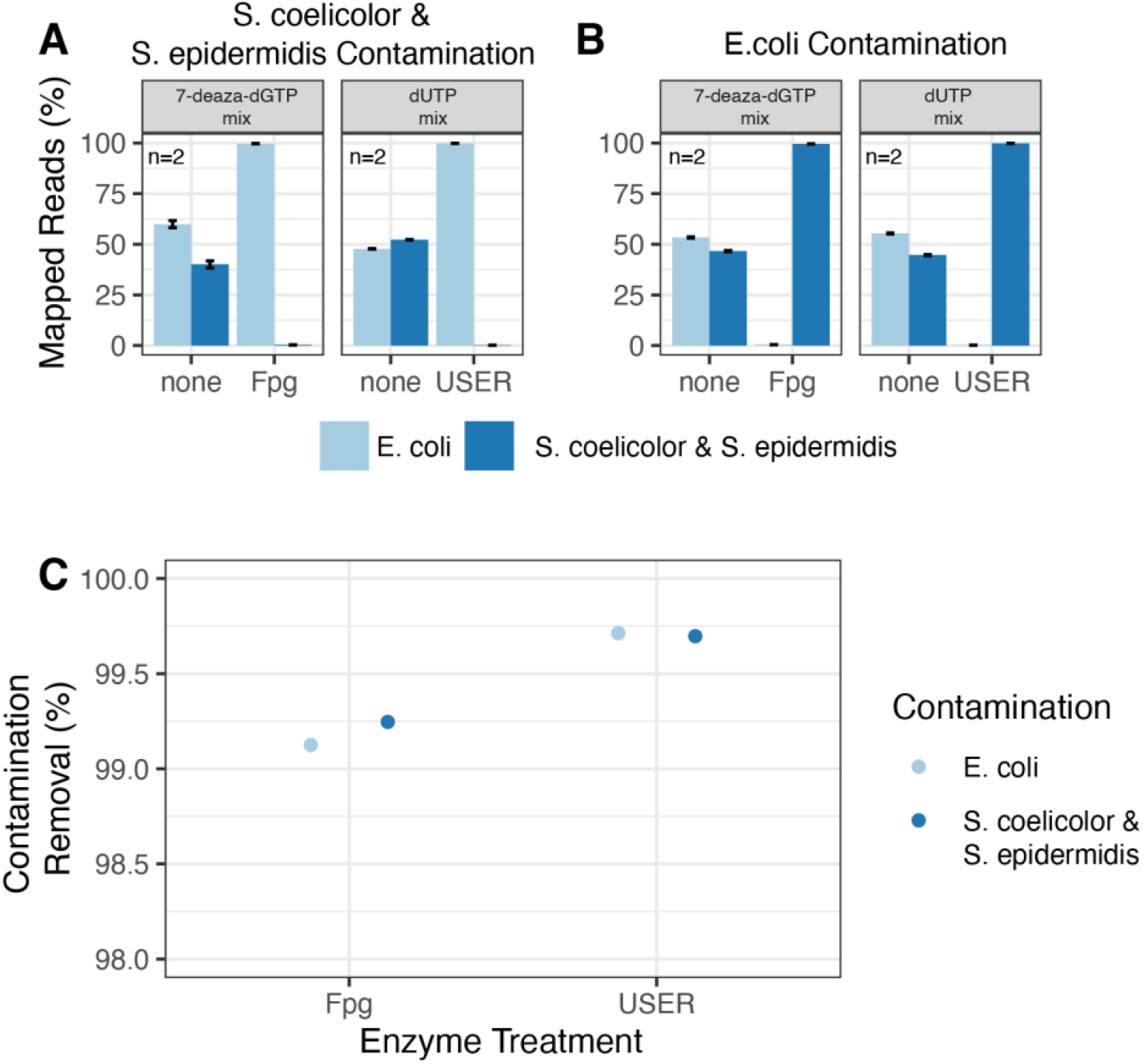
1:1 library contamination removal using Fpg or USER. E. coli and mixed S. coelicolor/S. epidermidis libraries were mixed at approximately 1:1 ratio, where one library has been amplified with dNTP mix, and the other had been amplified with either 7-deaza-dGTP mix or dUTP mix. Libraries were digested with Fpg, USER or no enzyme and again amplified with standard dNTPs before sequencing. Percent of reads mapping to the E. coli and S. coelicolor/S. epidermidis genomes with and without enzymatic treatment (Fpg or USER) is shown with the mean and standard deviation between the two replicates shown, where **A)** the S. coelicolor/S. epidermidis library is the contamination and **B)** the E. coli library is the contamination. **C)** Percent removal of carryover contamination after Fpg or USER treatment for S. coelicolor/S. epidermidis or E. coli contamination, calculated as (Cc – Ct)/Cc ^*^ 100, where Cc is the mean control reads (normalized by total mapped reads) from two replicates and Ct is the mean treatment reads (normalized by the total mapped reads) from two replicates.

To mimic a realistic PCR carryover contamination scenario, a pre-made *S. coelicolor/S. epidermidis* library amplified with either 7-deaza-dGTP mix or dUTP mix was spiked into *E. coli* genomic DNA at a 1:100 ratio. This setup simulates the introduction of residual amplicons from a prior amplification reaction into a fresh DNA sample. During library preparation, following adaptor ligation, these samples were treated with Fpg or USER and amplified using the appropriate nucleotide mix (7-deaza-dGTP mix or dUTP mix, respectively, **Figure 5A**). Untreated control libraries, processed in parallel under identical conditions but without enzymatic treatment, were used as references to evaluate the efficacy of carryover contamination removal. Approximately 0.01% of reads map back to the contamination after Fpg treatment as opposed to 2% in the control libraries. These values are comparable to USER treatment of dU-containing contamination **(Figure 5B**,**C)**. This corresponds to an estimated 96% and 98% removal in carryover contamination for Fpg and USER treatment respectively **(Figure 5D)**. Additionally, we tested the effect of different Fpg concentrations and incubation times on contamination removal in NGS libraries during the adaptor ligation library step. Overall, we did not observe a strong pattern, and most concentrations and incubation times resulted in similar levels of contamination removal **(Supplementary Figure 2)**.

**Figure 5:**
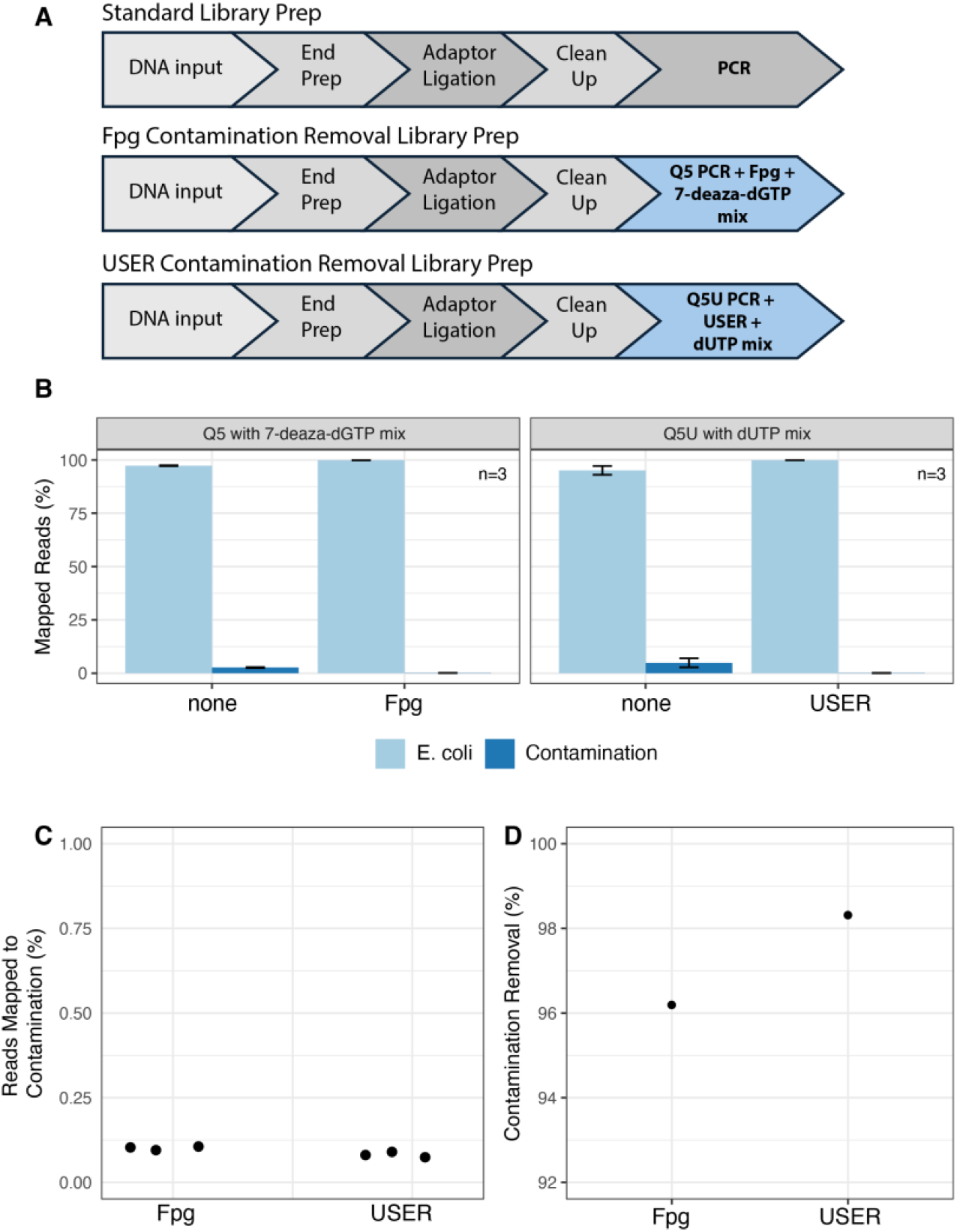
Removal of Amplicon Contamination Using Fpg Treatment. **A)** Schematic overview of the contamination library preparation workflow for USER or Fpg in comparison to the standard NEBNext Ultra II protocol. Contamination libraries were generated by spiking 0.1 ng of pre-amplified S. coelicolor/ S. epidermidis library—amplified with either a 7-deaza-dGTP mix (replacing dGTP) or a dUTP mix (replacing dTTP)—into 10 ng of E. coli genomic DNA. The contamination removal step (Fpg or USER treatment) was applied after adaptor ligation. **B)** Percent reads mapped with and without Fpg treatment (left) or USER treatment (right). **C)** Percent contamination with Fpg or USER treatment. **D)** Percent contamination removal with USER or Fpg treatment, calculated as (Cc – Ct)/Cc ^*^ 100, where Cc is the mean control reads (normalized by total mapped reads) from three replicates and Ct is the mean treatment reads (normalized by the total mapped reads) from three replicates.

**Supplementary Figure 2:**
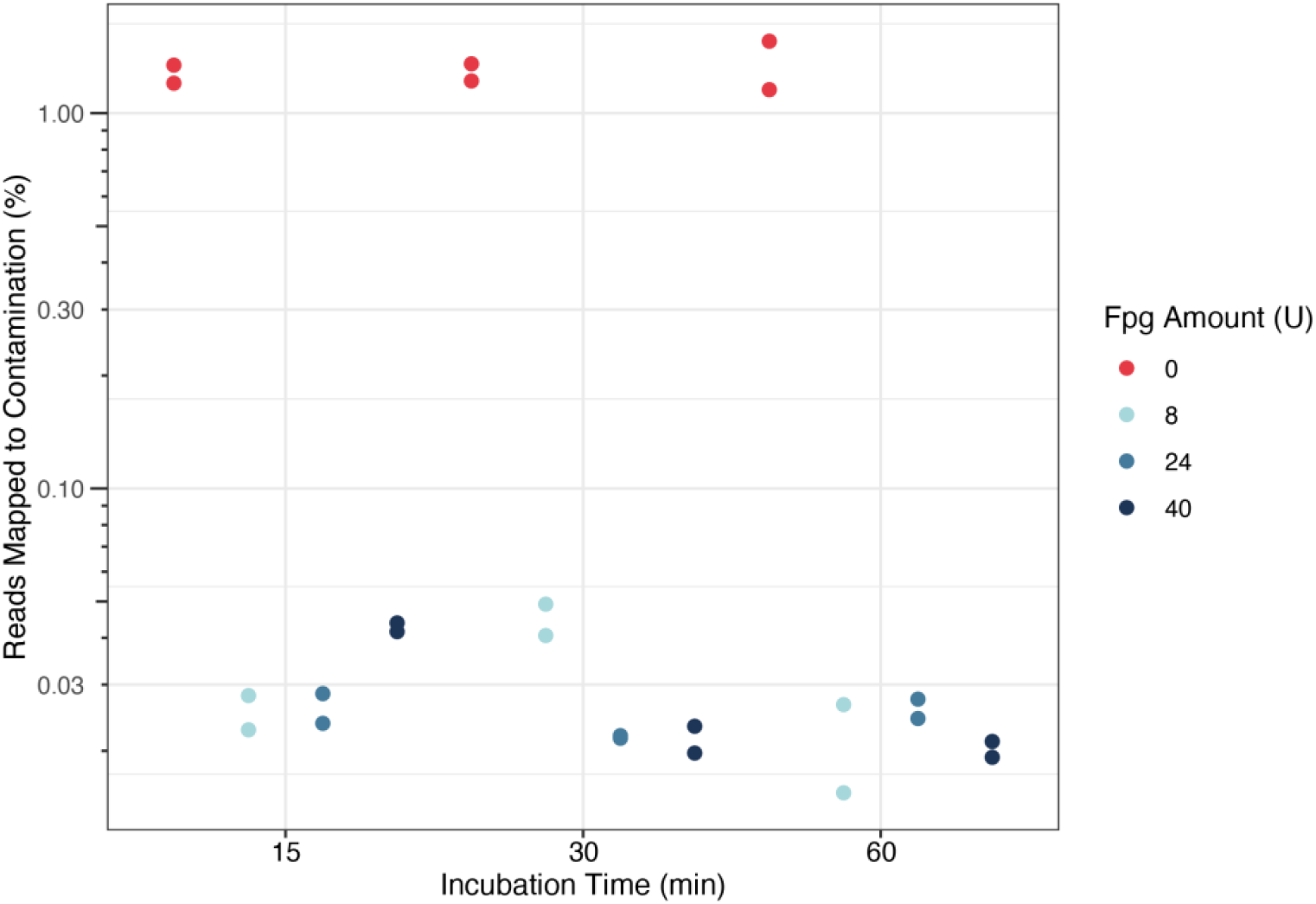
Fpg reaction time and concentration for contamination removal. 0.1ng of contamination DNA (amplified with 7-deaza-dGTP mix) was spiked into 10ng of E. coli DNA. Libraries were prepared in duplicate and treated with an increasing amount of Fpg (0, 8U, 24U, 40U) for 15, 30 or 60 minutes during the post adaptor ligation USER step. Libraries were amplified using Q5 with a 7-deaza-dGTP mix and percent reads mapping were calculated for each condition.

So far, all experiments have included a cleanup step following Fpg treatment to facilitate clearer data interpretation. However, for clinical applications, the Fpg treatment and subsequent amplification should ideally be performed in a single, closed-tube reaction to minimize the risk of contamination during the cleanup step. For this we tested Fpg concentrations and inactivation conditions directly within the PCR reaction mix with 7-deaza-dGTP fully replacing dGTP (referred to as Q5 PCR). Standard libraries were prepared, then treated with 0, 0.8, 4 or 8U of Fpg. For all conditions except the lowest amount of Fpg tested (0.8U), we observed significant depletion of library yield relative to no Fpg treatment, regardless of heat inactivation, while the library yield with the 0.8U of Fpg treatment is comparable to the control no Fpg condition **(Supplementary Figure 3)**. To optimize contamination removal at this lower Fpg concentration, libraries were prepared with 10ng *E. coli* spiked-in with 1ng of *S. coelicolor/S. epidermidis* 7-deaza-dGTP-amplified contamination and treated with either 0.8U (0.1 ul) of Fpg or no treatment (schematic in **Figure 6A**) and incubated at 0, 15, 30, or 60 min at 37°C before proceeding into PCR amplification. We again observe greater than 97% contamination removal regardless of incubation time at 37°C, with less than 0.25% of reads mapping back to the contamination sample **(Figure 6 B**,**C**,**D)**. However, incubation for longer than 15 minutes at 37°C prior to PCR reduces library yield **(Supplementary Figure 4)**.

These results demonstrate that a low concentration of Fpg in the PCR mix is highly effective at eliminating carryover contamination. Furthermore, only a brief pre-amplification incubation confers full carryover contamination prevention for NGS library preparation without compromising library yield.

Additionally, we tested a condition in which 7-deaza-dGTP was included in a Q5 master mix (referred to as Q5 MM) already containing dNTPs, resulting in a 1:1 ratio of dGTP to 7-deaza-dGTP. Under these conditions, we still observe only 0.5% reads mapping back to the contamination compared to 10.4% reads in the non-treated sample corresponding to more than 92% contamination removal **(Supplementary Figure 5)**. Q5 PCR and Q5 MM libraries were checked for GC bias and error rate distribution. Interestingly, libraries treated with Fpg during PCR have significantly lower G to T (read 2) and C to A (read 1) substitutions, suggesting Fpg can additionally repair low levels of oxidative damage prior to PCR **(Figure 6E, Supplementary Figure 6)**.

Overall, this method successfully removes contamination with the additional benefit of reducing GC biases and flexibility in polymerase utilization for NGS applications. Additionally, the workflow does not require additional steps and can be easily incorporated in standard NGS workflows.

**Supplementary Figure 3:**
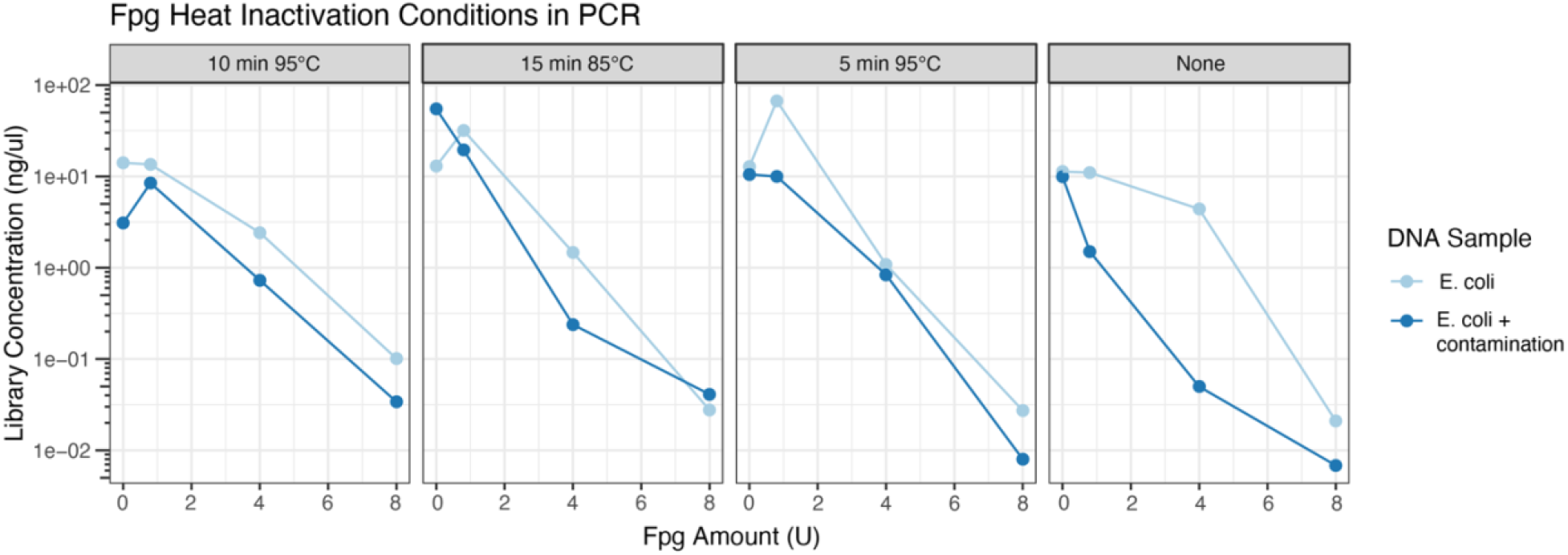
Library yield is inversely correlated to Fpg concentration in the PCR mix. Libraries were prepared for E. coli genomic DNA (light blue) or a mix of E. coli genomic DNA and a contamination of S. coelicolor/S. epidermidis library amplified with the 7-deaza-dGTP mix (dark blue). Prior to the amplification step various amounts of Fpg (0, 0.8, 4 or 8U) and heat inactivation conditions were tested (None, 5 minutes at 95°C, 10 min at 95°C or 15 min at 85°C). Library yields were quantified by Agilent TapeStation HSD1000 reagents for each sample.

**Figure 6:**
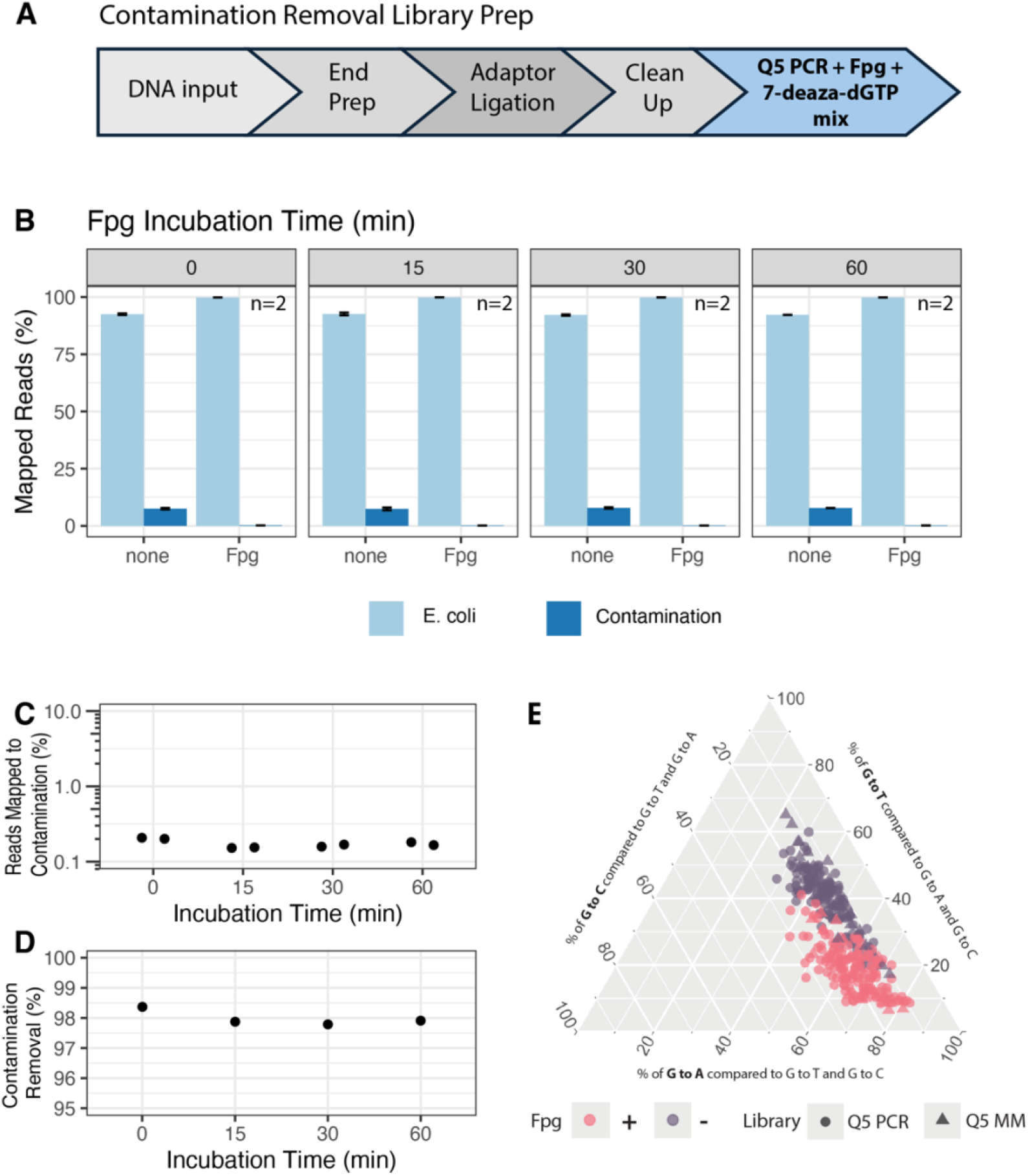
Low input Fpg and short incubation time is sufficient for contamination removal during PCR. **A)** Schematic overview of the contamination removal workflow in which Fpg (0.1 µL, 0.8 U) and 7-deaza-dGTP (fully substituting for dGTP) are incorporated directly into the final Q5 PCR amplification step to enable complete carryover protection. Libraries were prepared using a 1:10 ratio of contamination (S. coelicolor/S. epidermidis library amplified with 7-deaza-dGTP) to target DNA (E. coli genomic DNA). **B)** Percent reads mapped with and without Fpg after incubation at 37°C pre-PCR for 0, 15, 30 or 60 minutes. **C)** Percent contamination reads under each incubation time. **D)** Percent contamination removal after each incubation time, calculated as (Cc – Ct)/Cc ^*^ 100, where Cc is the mean control reads (normalized by total mapped reads) from two replicates and Ct if the mean treatment reads (normalized by the total mapped reads) from two replicates. **E)** Ternary plot showing the relative contribution of G to T compared G to A and G to C substitutions in libraries prepared with or without Fpg in a Q5 PCR mix with 7-deaza-dGTP fully replacing dGTP (Q5 PCR) or with 7-deaza-dGTP added into the premade Q5 master mix at 1:1 ratio with dGTP (Q5 MM).

**Supplementary Figure 4:**
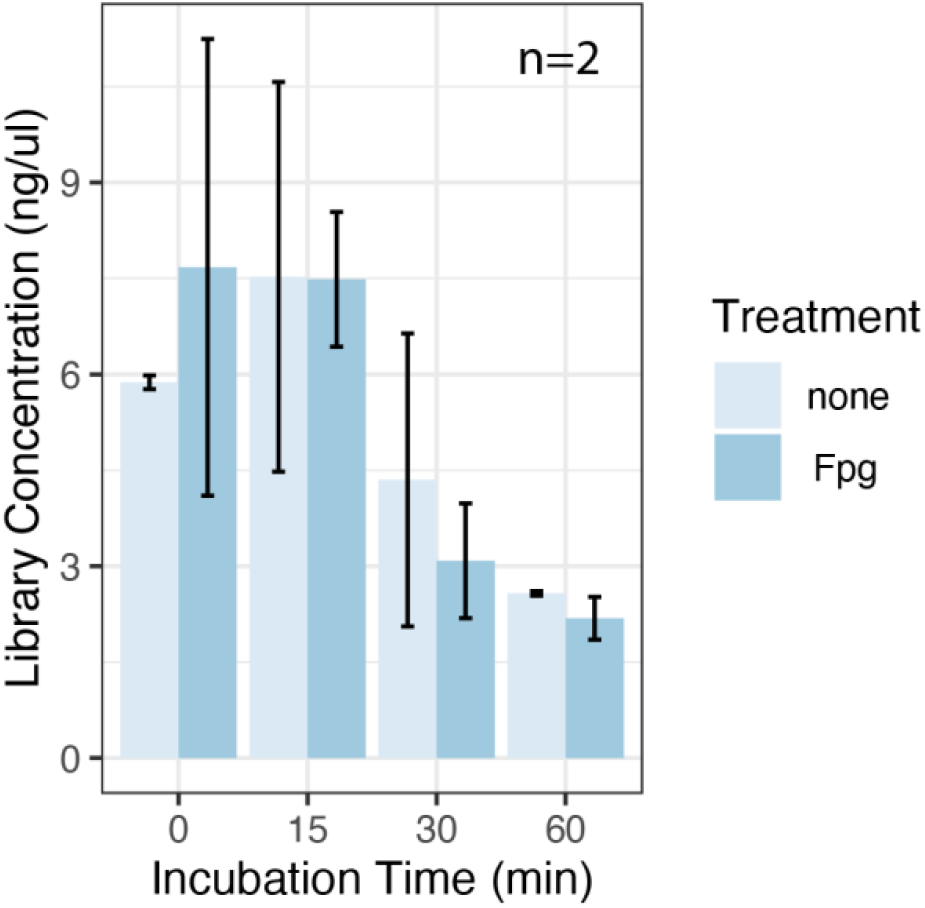
Library yield after Fpg treatment within Q5 PCR: Libraries were prepared using a 1:10 ratio of contaminant to target DNA in which Fpg (0.1 µL, 0.8 U) and 7-deaza-dGTP (fully substituting for dGTP) were incorporated directly into the final Q5 PCR amplification step to enable complete carryover protection. Fpg treatment during PCR was tested at 0, 15, 30, and 60 min. Library yield post amplification for each condition quantified by Agilent TapeStation HSD1000 reagents for each condition.

**Supplementary Figure 5:**
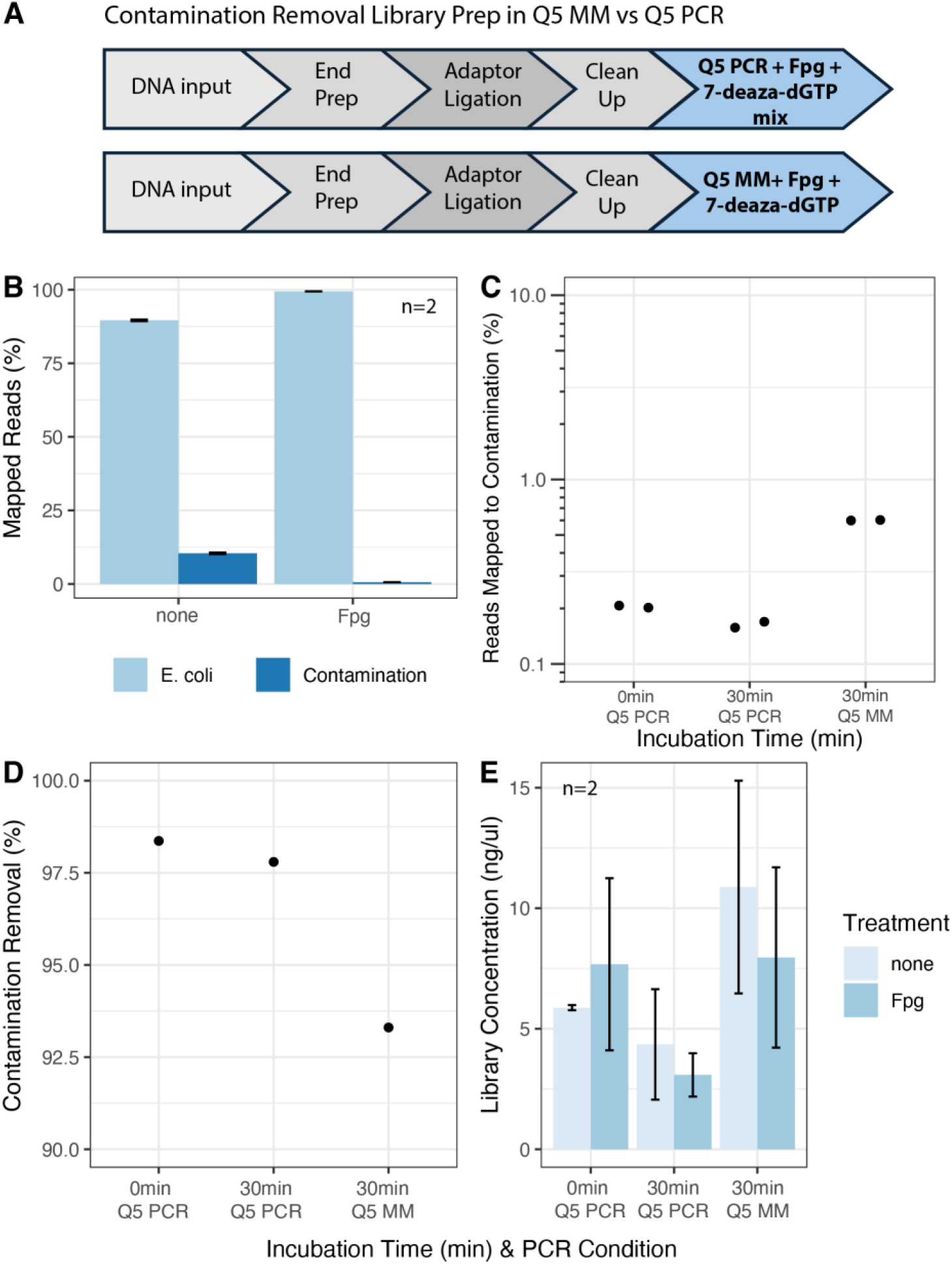
Low input Fpg and supplemented 7-deaza-dGTP reduce carryover contamination in NGS Q5 Master Mix. **A)** Contamination removal workflow where Fpg (0.1ul, 0.8U) and 7-deaza-dGTP are included in the PCR step for complete carryover protection either fully replacing dGTP in a custom reaction mix with Q5 polymerase (Q5 PCR, top) or spiked into Q5 Master Mix (Q5 MM, bottom) at 1:1 ratio to dGTP. Libraries were prepared using a 1:10 ratio of contamination (S. coelicolor/S. epidermidis library amplified with 7-deaza-dGTP) to target DNA (E. coli genomic DNA). **B)** Percent reads mapped for Q5 MM sample incubated for 30 min. at 37°C with or without Fpg. **C)** Percent contamination reads for Q5 PCR incubated for 0 or 30 min and Q5 MM incubated for 30 min. **D)** Percent contamination removal for Q5 PCR and Q5 MM, calculated as (Cc – Ct)/Cc ^*^ 100, where Cc is the mean control reads (normalized by total mapped reads) from two replicates and Ct is the mean treatment reads (normalized by the total mapped reads) from two replicates. **E)** Library yield post amplification for each condition quantified by Agilent TapeStation HSD1000 reagents for each condition.

**Supplementary Figure 6:**
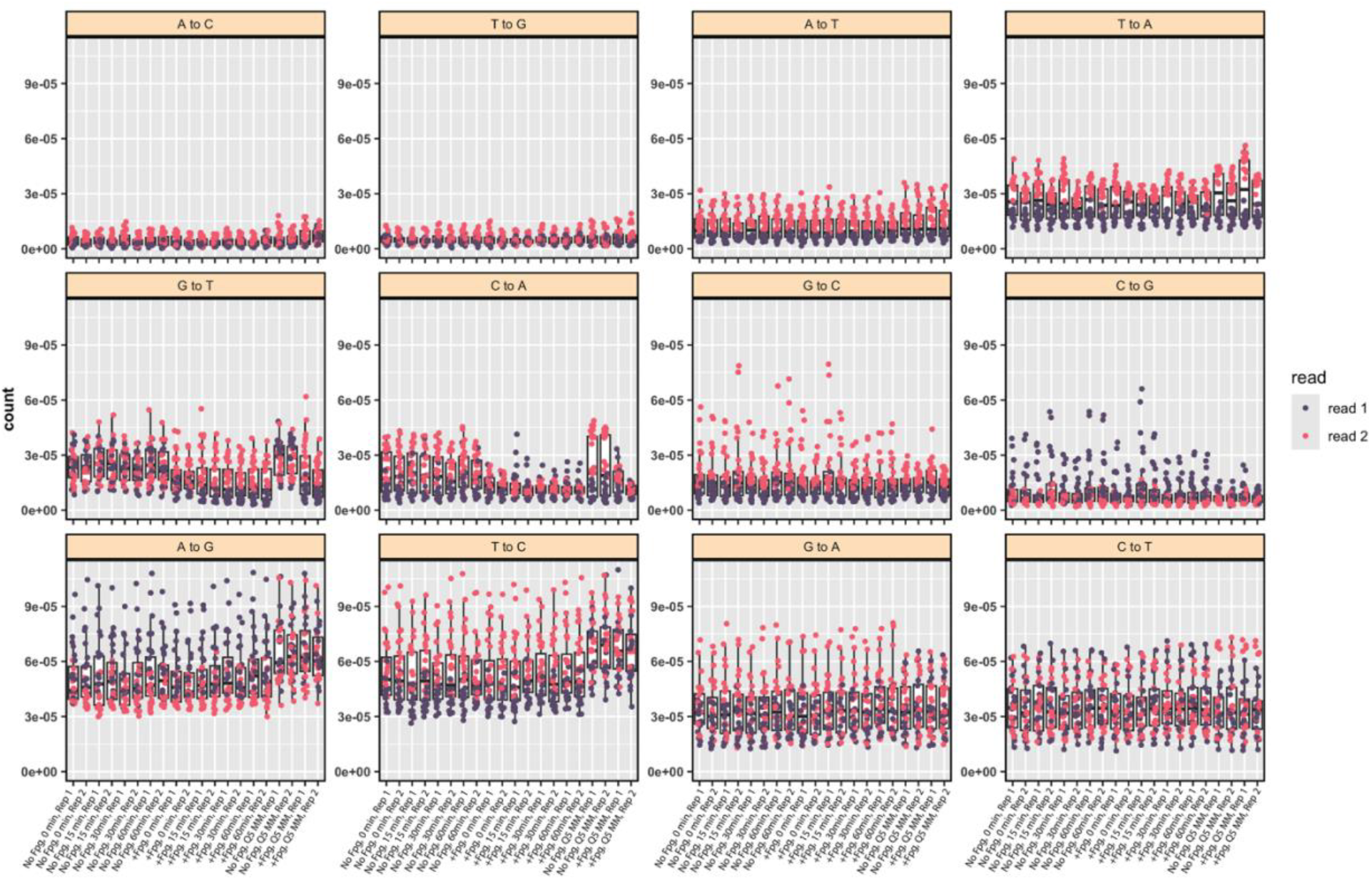
Error rates in NGS data from contamination experiment with Fpg. For each substitution type, frequencies were calculated separately for Read 1 (purple) and Read 2 (pink) across all 5′NXN3′ sequence contexts (where N = A, T, C, or G, and X denotes one of the 12 substitution types). Data are presented for libraries amplified with 7-deaza-dGTP mix and treated with or without 0.8U of Fpg and amplified with either a Q5 PCR mix where 7-deaza-dGTP fully replaces dGTP or in which 7-deaza-dGTP was spiked in a Q5 master mix at 1:1 ratio with dGTP (Q5 MM).

## Discussion

The use of dUTP incorporation during PCR amplification, followed by UDG treatment, is a well-established method for preventing carryover contamination, particularly in diagnostic applications. However, this approach requires polymerases engineered to tolerate uracil residues without stalling, limiting its broader applicability. In this study, we present a novel and mechanistically distinct approach using the Fpg/7-deaza-dGTP pair. We demonstrate that this method is compatible with standard DNA polymerases, thereby expanding its utility to next-generation sequencing workflows and other amplification-based applications for which specialized enzymes capable of efficiently bypassing uracil residues are not desired.

We show that NGS libraries amplified with 7-deaza-dGTP have sequencing quality that match or exceed standard library preparation procedures. Additionally, 7-deaza-dGTP has the additional benefit of a more even genome wide coverage, particularly across GC and AT rich regions where polymerases generally are impaired. This simple method requires reagents that are commercially available, can be easily incorporated into standard NGS workflows and does not detrimentally impact sequencing results.

Importantly, substitution frequency is generally not impaired when 7-deaza-dGTP is incorporated, unlike other guanosine analogs such as inosine and 8-oxo-guanosine^16^. Additionally, we even observe a slightly lower G to T substitution frequency for 7-deaza-dGTP. This may suggest that 7-deaza-deoxyguanosine confers lower in situ oxidative damage during library prep and sequencing compared to standard deoxyguanosine. Additionally, when Fpg is included for carryover, we observed further decreased G to T substitution frequency. While these libraries were sheared with Covaris in 1X TE buffer, residual oxidative damage may have occurred^17^, and incorporating Fpg can repair this damage within the PCR step. We also demonstrate that this approach can be seamlessly integrated into the library preparation workflow without adding time or steps, by including Fpg directly in the PCR amplification mix.

One drawback of this approach is quantification of the DNA after amplification with 7-deaza-dGTP. Although one of its main advantages is reducing Hoogsteen bonding in GC base pairs, this also impacts intercalating dyes such as ethidium bromide and Qubit quantification methods^18,19^. As a result, quantification methods relying on these dyes must be specifically adjusted to account for the presence of 7-deaza-dGTP. Alternatively, using a mix of these methods as well as spectrophotometric readings such as Nanodrop or Lunatic (Unchained Labs), or methods like TapeStation be used to quantify fully modified 7-deaza-deoxyguanosine-containing amplicons.

In summary, this novel 7-deaza-dGTP/Fpg strategies for carryover prevention can be important for many applications from forensics, molecular diagnostics and low input or precious sample sequencing such as single cell sequencing, environmental samples and ancient DNA. While not yet tested, this method may also work well in qPCR, standard PCR and LAMP contamination removal assays in place of UDG/dUTP

## Material and Methods

### 4.1 Digestion of 7-deaza-deoxyguanosine in DNA

Primers were designed from the λ (Lambda) genome (Lambda DNA, NEB #M0481) and synthesized by IDT to amplify 300 bp (F: 5’TGCCTGCAAAGATGAGGAGG3’ and R: 5’GCATTCACCGCACCGATAAC3’) and 1200 bp fragments (F: 5’GAGTGGCGGCAGATAAAGGT3’ and R: 5’CCCCGCCAGATGATAAGCAT3’). PCR amplification was performed using Q5 Hot Start High-Fidelity DNA Polymerase (NEB #M0493) with two different nucleotide mixes: (1) a standard dNTP mix to generate a canonical 1.2 kb amplicon, and (2) a modified mix containing dATP, dTTP, dCTP, and 7-deaza-dGTP (NEB #N0445) to produce a fully modified 300 bp amplicon in which all deoxyguanosine are substituted with 7-deaza-deoxyguanosine. Both amplicons were quantified by TapeStation genomic DNA reagents (Agilent #5067-5365) and mixed before treatment with either Fpg (NEB #M0240S, 8U) or no enzyme for 30 minutes at 37°C in rCutsmart buffer (NEB #B6004). After digestion and purification, samples were run on a 1.2% Agarose gel together with 1 kb Plus DNA Ladder (NEB #N3200) to visualize the digestion of amplicons.

### 4.2 Library Construction

Two separate libraries were constructed from [1] Genomic DNA from *Escherichia coli* K-12 strain DHB4 (CP014270) and [2] a mix of genomic DNA from *Streptomyces coelicolor* (ATCC #10147) and *Staphylococcus epidermidis*. For this, genomic DNA was sheared to 250bp (20ng/ul DNA, 1X TE 50ul, Covaris AFA minitubes (Covaris PN 500514), 10% intensity, 5 cycles, 200 Burst, 40seconds). Using 100ng of sheared gDNA per sample, NEBNext Ultra II DNA Library Preparation (NEB #E7645) reagents were used for end preparation repair and adaptor ligation following manufacturer recommendations. Unamplified libraries were named [1] *E*.*coli* unamplified library and [2] *S. coelicolor/S*.*epidermidis* unamplified library.

Both libraries were PCR amplified with each of the following master mixes: 1) Q5 (NEB #M0493) with standard 0.2mM dNTPs, 2) Q5 with 0.2mM 7-deaza-dGTP mix (d[ATC]TP + 7-deaza-dGTP) and 3) Q5U (NEB #M0515) with 0.2mM dUTP mix (d[ACG]TP with dUTP). NEBNext Unique Dual Indices (NEB #E6440S) for Illumina at recommended concentrations were used for amplification. The resulting 6 amplified libraries were named *E*.*coli_*dNTP, *E*.*coli_*7-deaza-dGTP, *E*.*coli_*dUTP, *S*.*coelicolor/S*.*epidermidis_dNTP, S*.*coelicolor/S*.*epidermidis_*7-deaza-dGTP and *S*.*coelicolor/S*.*epidermidis_*dUTP. Remaining amplification conditions and cycling were done following manufacturer recommendations.

These amplified libraries *E*.*coli_*dNTP, *E*.*coli_*7-deaza-dGTP, *E*.*coli_*dUTP, *S*.*coelicolor/S*.*epidermidis_dNTP, S*.*coelicolor/S*.*epidermidis_*7-deaza-dGTP and *S*.*coelicolor/S*.*epidermidis_*dUTP were sequenced on Illumina NovaSeq 6000 SP paired-end reads with an average of 200 million reads per sample.

### 4.3 Mixed Libraries

Using amplified libraries prepared as described above, *E. coli* and *S. coelicolor* libraries were combined at a 1:1 ratio (*E*.*coli*_dNTP + *S*.*coelicolor/S*.*epidermidis_*dUTP, *E*.*coli*_dUTP + *S*.*coelicolor/S*.*epidermidis_*dNTP, *E*.*coli*_dNTP + *S*.*coelicolor/S*.*epidermidis_*7-deaza-dGTP, *E*.*coli_*7-deaza-dGTP + *S*.*coelicolor/S*.*epidermidis_*dNTP). Samples were treated with 1ul of either USER (1U/ul, NEB #M5505), Fpg (8U/ul, NEB #M0240), or nuclease-free H_2_O (no enzyme control) at 37°C for 15 min in rCutsmart Buffer and then cleaned with NEBNext Sample Purification Beads following NEBNext guidelines for a 1X bead purification. Samples were amplified as described above with Q5 Ultra II MM or Q5U MM for dUTP containin libraries with NEBNext Unique Dual Indices. Samples were sequenced on NextSeq 500 Mid Output with paired-end reads.

### 4.4 Simulated contamination libraries

To test the removal of 7-deaza-dGTP contamination, 0.1ng of *S*.*coelicolor/S*.*epidermidis_*7-deaza-dGTP or *S*.*coelicolor/S*.*epidermidis_*dUTP library prepared above was spiked into 10ng of *E. coli* genomic DNA. End repair and adapter ligation steps were followed according to manufacturer recommendations. During the USER treatment step, after adaptor ligation, 3ul (24U) of Fpg or 3ul nuclease-free H_2_O (no enzyme control) was added and incubated at 37°C for 15 minutes in triplicate. Samples were bead purified following NEBNext manual recommendations, then amplified with Q5 and standard dNTPs, Q5 with 7-deaza-dGTP mix, or Q5U with dUTP mix with NEBNext Unique Dual Indices as above. Samples were sequenced on Illumina NovaSeq 6000 SP paired-end reads.

### 4.5 Fpg Digestion Conditions

To test the impact of time and Fpg amount on contamination removal, contamination libraries were prepared with 0.1ng of *S*.*coelicolor/S*.*epidermidis_*7-deaza-dGTP library spiked into 10ng of *E. coli* genomic DNA. After adaptor ligation, 0, 8, 24 or 40U of Fpg (8U/ul) was added to the reaction and incubated at 37°C for 15, 30 or 60 min and prepared in duplicate. Libraries were amplified with Q5 with 7-deaza-dGTP mix and sequenced with paired-end reads on Illumina NovaSeq 6000 SP.

Next, Fpg was tested directly in the PCR mix during library preparation. Contamination libraries were prepared with 1ng of *S*.*coelicolor/S*.*epidermidis_*7-deaza-dGTP spiked into 10ng of *E. coli* genomic DNA, following NEBNext Ultra II manufacturer recommendations until the PCR step. Q5 PCR mixes were prepared in duplicate with 7-deaza-dGTP with 0, 0.8, 4 or 8U of Fpg (8U/ul) in a 50ul reaction. Four Fpg heat inactivation conditions were tested: none, 5 minutes at 95°C, 10 minutes at 95°C or 15 min at 85°C, then immediately proceeding into the PCR. After amplification, libraries were bead purified at 0.9X with NEBNext Sample Purification beads and run on an Aglient TapeStation HSD1000 for quantification and visualization.

After determining optimal Fpg input and heat inactivation conditions within the PCR amplification, libraries were prepared to determine if lower Fpg amounts within a PCR produced adequate carryover prevention. As above, contamination libraries were prepared with 1ng of *S*.*coelicolor/S*.*epidermidis_*7-deaza-dGTP spiked into 10ng of *E. coli* genomic DNA. Libraries were then amplified in duplicate with the following conditions: 0 or 0.8U (0.1ul) of Fpg and incubated at 37°C for 0, 15, 30 or 60 minutes before directly proceeding to the PCR and amplified with 7-deaza-dGTP Q5 mix. Additionally, libraries were amplified with standard Q5 MM supplemented with 7-deaza-dGTP (final concentration: 0.2mM) with 0 or 0.8U of Fpg and incubated at 37°C for 30 min, before proceeding to the PCR. Libraries were sequenced on NovaSeq SP 6000 paired-end reads.

### 4.6 Bioinformatic analysis

For GC bias analysis and error rate estimation, paired-end reads were mapped to Staphylococcus epidermidis strain SESURV_p1_0557 (CP043777.1), *Streptomyces coelicolor* (ATCC #10147) and *Escherichia coli* K-12 strain DHB4 (CP014270.1) using bowtie2. GC bias and fragment size distributions were calculated using Picard (Version:2.26.4) CollectGcBiasMetric and CollectInsertSizeMetrics respectively. The error rate estimation was performed using the damage estimator software (https://github.com/Ettwiller/Damage-estimator) with split_mapped_reads.pl and estimate_damage_context.pl (default parameters).

For the contamination analysis, reads were trimmed with trimgalore (default parameters, version 0.6.10), then mapped to a combined reference genome of *Streptomyces coelicolor* (ATCC #10147), *Staphylococcus epidermidis* strain SESURV_p1_0557 (CP043777.1), and *Escherichia coli* K-12 strain DHB4 (CP014270.1) using bwa mem (version 0.7.17-r1188). Mapped reads were filtered to exclude low quality mapping and multiple mapping using samtools (-q 60 flag, version 1.17), and the reads mapped per genome were extracted with samtools idxstats. Percent mapping to each genome was calculated as (reads per genome per sample / total mapped reads per sample)^*^100. Contamination removal was calculated as (C_C_ – C_T_)/C_C_ ^*^ 100, where C_C_ is the mean of the normalized control reads, and Ct is the mean normalized treatment reads (Fpg or USER treated). Control reads and treatment reads were normalized by the total number of mapped reads for each sample.

## Acknowledgements

The authors gratefully acknowledge the following for providing materials, services, and support during this project: NEB Sequencing Core and NEB IT Department for technical services. Lastly, the authors are very grateful to the Comb family and New England Biolabs, Inc. for funding the entirety of this work.

## Data and code availability

All raw and processed sequencing data generated in this study have been submitted to the NCBI Sequence Read Archive (SRA; https://www.ncbi.nlm.nih.gov/geo/) and will be made public upon publication.

